# Transcription Factor RFX3 Stabilizes Mammary Basal Cell Identity

**DOI:** 10.1101/2021.12.13.472491

**Authors:** Erica M. Tross, Christian R. de Caestecker, Ken S. Lau, Ian G. Macara

## Abstract

The myoepithelial cell compartment of the murine postnatal mammary gland is generated from basal cap cells in the terminal end bud and maintained by self-renewal. Transdifferentiation to the luminal lineage does not normally occur but can be induced by DNA damage, luminal cell death or transplantation into a recipient mammary fat pad. Myoepithelial cells cultivated in vitro can also transdifferentiate towards the luminal lineage. Little is known about the molecular mechanisms and gene regulatory networks underlying this plasticity. Using a transgenic mouse (Tg11.5kb-GFP) that marks cap cells with GFP, we discovered that mature myoepithelial cells placed in culture begin to express GFP within ∼24 hrs and later express the Keratin 8 (K8) luminal marker. Cell tracking showed that most K8+ cells arose from GFP+ cells, suggesting that myoepithelial cells de-differentiate towards a progenitor state before changing lineage. Differential gene expression analysis, comparing pure GFP+ cap cells with mature myoepithelial cells, identified multiple transcription factors that iRegulon predicted might regulate the myoepithelial to cap cell transition. Knockout of one of these genes, Regulatory Factor 3 (Rfx3), significantly reduced the population of GFP+ cells and increased differentiation to the K8+ luminal lineage. Rfx3 knockout also reduced mammosphere growth and mammary gland regeneration efficiency in a transplantation assay, but had no effect on proliferation in vitro. Together, these data support a key role for Rfx3 in the stabilization of the mammary basal cell lineages.

## INTRODUCTION

The mammary gland undergoes dynamic changes that contribute to the development, structure and function of the organ. In utero, development is initiated by bipotent mammary stem cells that give rise to the mammary rudiment, which remains quiescent until puberty (Spike et al., 2012; Van Keymeulen et al., 2011; Wuidart et al., 2018). At the onset of puberty, the estrus cycle drives expansion of the rudiment into a branching network of ducts that fill the mammary fat pad (Silberstein, 2001). Mature ducts consist of an inner layer of polarized luminal cells (which express the keratin K8), surrounded by spindle-shaped basal, myoepithelial cells that are marked by K14 expression. Outgrowth occurs primarily from terminal end buds at the tip of each duct, filled with immature, proliferating luminal body cells enveloped by a single basal layer of cap cells, which are a metastable population of myoepithelial progenitors (Abdul-Manan et al., 1999; Humphreys et al., 1996; Sreekumar et al., 2017). These large end buds, and the cap cells, disappear when the ducts reach the edges of the fat pad (Hinck & Silberstein, 2005). At homeostasis the basal and luminal lineages are lineage-restricted and maintained separately by self-renewal (Prater et al., 2014; Van Keymeulen et al., 2011; Wuidart et al., 2016). Each population appears to be stable; however, we discovered that DNA damage to the mammary gland triggers transdifferentiation of myoepithelial to luminal cells (Seldin and Macara 2020). Ablation of luminal cells in situ by diphtheria toxin also activates transdifferentiation, possibly through multiple signaling pathways (Centonze et al., 2020; Le Guelte, 2021). Interestingly, this same lineage switch appears to be induced by cultivating myoepithelial cells in vitro or by their transplantation into cleared mammary fat pads in a recipient mouse. However, almost nothing is known about the underlying mechanisms and gene regulatory networks that determine the stability of the lineages or their reversion to a stem-like state. To begin to untangle the processes driving myoepithelial cell transdifferentiation, we employed a transgenic mouse strain that expresses GFP from the promoter of the s-SHIP gene (11.5kb-GFP), which is expressed in embryonic and hematopoietic stem cells, and several other progenitor/stem cell lineages (Rohrschneider, Custodio, Anderson, Miller, & Gu, 2005). It also marks cap cells but is silent in mature myoepithelial cells (Bai & Rohrschneider, 2010). After plating purified myoepithelial cells in culture, we detected both GFP+ cells and cells expressing the luminal marker K8 arising within 24 – 48 hrs. Single cell RNAseq also identified a population of luminal cells after 96 hrs culture. Cell tracking revealed that most K8+ cells arise from the GFP+ population, suggesting that myoepithelial cells de-differentiate before undergoing transdifferentiation towards the luminal lineage. Bulk RNAseq of purified cap cells (GFP+) and mature myoepithelial cells identified several transcription factors that were preferentially expressed in the cap cell population, which by iRegulon analysis are predicted to control multiple other genes induced in the cap cells. For one such factor, Rfx3 (regulatory factor 3), its ablation by CRISPR-Cas9 editing in mature myoepithelial cells caused a significant reduction in the frequency of GFP+ cell formation, with increased formation of K8+ cells. Rfx3 knockout also reduced mammosphere formation and reduced ductal outgrowth after transplantation. Rfx3 ablation in pure cap cells also increased the formation of K8+ cells. We propose that Rfx3 normally stabilizes basal cell identities, so that in its absence the myoepithelial and cap cell populations more rapidly convert to the luminal lineage, thereby losing the ability to regenerate mammary glands in transplantation assays.

## RESULTS

### Myoepithelial cells in culture can transdifferentiate towards cap cell and luminal cell lineages

Lineage-restricted mammary myoepithelial cells in situ can transdifferentiate into luminal cells in response to DNA damage or DTA-mediated ablation of the luminal population, and are capable of regenerating entire mammary glands after transplantation into the fat pads of recipient mice (Bai & Rohrschneider, 2010; Centonze et al., 2020; Seldin & Macara, 2020; Stingl et al., 2006; Van Keymeulen et al., 2011). To probe mechanisms that might regulate this lineage switching, we isolated mammary glands from the 11.5kb-GFP transgenic mouse, which expresses GFP specifically in cap cells of terminal end buds but not in mature myoepithelial cells (Supplementary Fig S1 A). Myoepithelial cells were then sorted by FACS (Supplementary Figure S1 B) and cultured for 96 hrs on laminin-coated cover glasses in FAD media supplemented with ROCK inhibitor (Fig 1 A). Cell cultures were fixed every 24 hrs over 4 d, stained for the luminal marker K8 and imaged to determine whether cells expressed GFP and/or K8. GFP+ cells began to appear after 24hrs and increased over time. K8+ cells were detectable within 48 hrs. By 96 hrs in vitro ∼20% of cells were GFP+, ∼10% of cells expressed K8 and ∼5% of cells were double positive (Fig 1 B,C).

**Figure 1.**
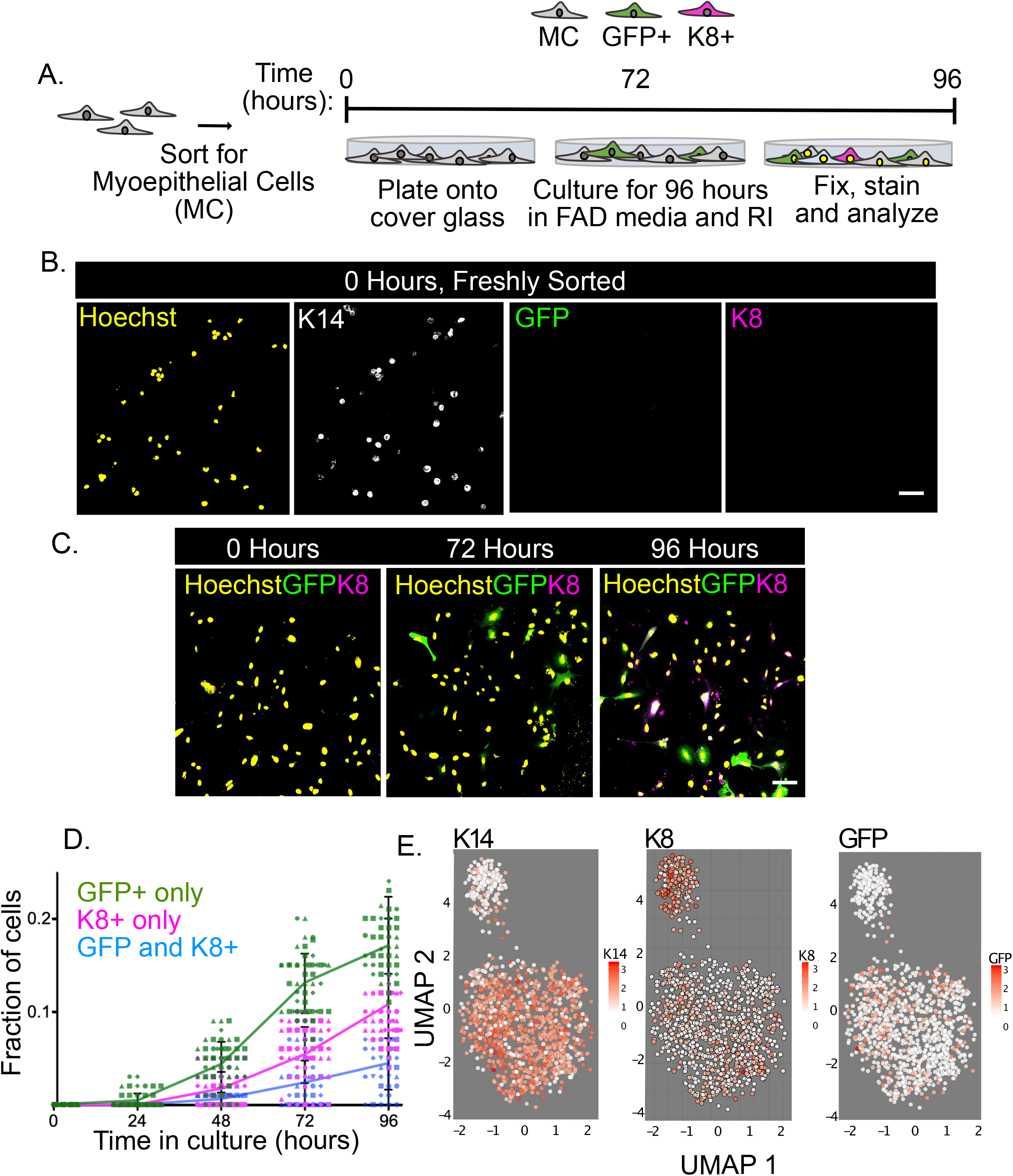
Myoepithelial cells from transgenic mice (TG11.5kb-GFP) express GFP and luminal marker, K8, after short-term culture. A. Diagram of experimental approach used to follow transdifferentiation of myoepithelial cells (MC) into luminal marker-expressing K8+ cells. B. Representative image of freshly sorted myoepithelial cells at 0 hrs in culture, expressing only myoepithelial cell marker K14. Scale bar = 100 µm C. Representative images of myoepithelial cells cultivated in FAD medium with ROCK inhibitor Y-27632 (RI) over 96 hrs. Scale bar = 100 µm D. Quantification of cap cell and luminal cell marker expression during culture of myoepithelial cells with Y-27632. A fraction of cells become GFP+, K8+ or express both markers, n= 5 biological replicates (each a different shape), 10 fields/experiment. Error bars = Mean +/- SD. E. UMAP projections of myoepithelial cells after 96 hrs culture. Single Cell RNA sequencing was used to observe myoepithelial cell gene expression changes over 96 hours.

To determine if this K8+ population represents the true luminal lineage or is a de-differentiated cell type that happens to express this keratin but not other luminal markers, we performed single cell RNAseq (scRNAseq) (Supplementary Fig S1 C). As an initial test we purified myoepithelial, luminal and cap cells from Tg11.5kb-GFP mouse mammary glands, then pooled equal numbers of each population and processed them for inDrop scRNAseq. Uniform manifold approximation and projection (UMAP) identifies three separate clusters, as expected (Supplementary Fig S1 D). We then isolated mature myoepithelial cells and grew them in culture for 96 hrs prior to encapsulation and scRNAseq. In this case UMAP revealed a large cluster that expresses the myoepithelial marker K14 and a separate cluster mostly positive for K8 and other luminal markers, while GFP+ cells were clustered with the K14+ population (Fig 1 D). These data suggest that the GFP+ cells do not fully de-differentiate into cap cells, but that transdifferentiation into the luminal lineage is relatively robust. We next asked if the GFP+ state is an obligatory intermediate between the myoepithelial and luminal identities, or whether different myoepithelial cells independently adopt either the luminal state or the GFP+ state. To distinguish these hypotheses, we used cell tracking. Freshly isolated myoepithelial cells from Tg11.5kb-GFP mice were plated sparsely on gridded coverslips and imaged every 12 hrs for 96 hrs then fixed and stained for K8 (Fig 2 A,B). Each cell was then traced backwards in time to determine if/when cells became GFP+ and if they progressed or not to become K8+. Data were visualized as a Sankey plot (Fig 2 C). Of 402 cells that were analyzed, 70% began to express GFP within 72 hrs and of this population ∼85% eventually expressed K8. In contrast, only 30% of the GFP-cells eventually expressed K8 (Fig 2 C). It is possible that some of these cells transiently expressed GFP and were missed during the periodic imaging. Moreover, we noted that transdifferentiation of this sparse culture of myoepithelial cells was much more efficient than occurs when the cells are plated more densely, as in Fig 1. The reason for this difference remains obscure but we can conclude, nonetheless, that the principal pathway for transdifferentiation involves an initial, partial dedifferentiation towards a progenitor, cap cell state prior to converting to the luminal lineage.

**Figure 2.**
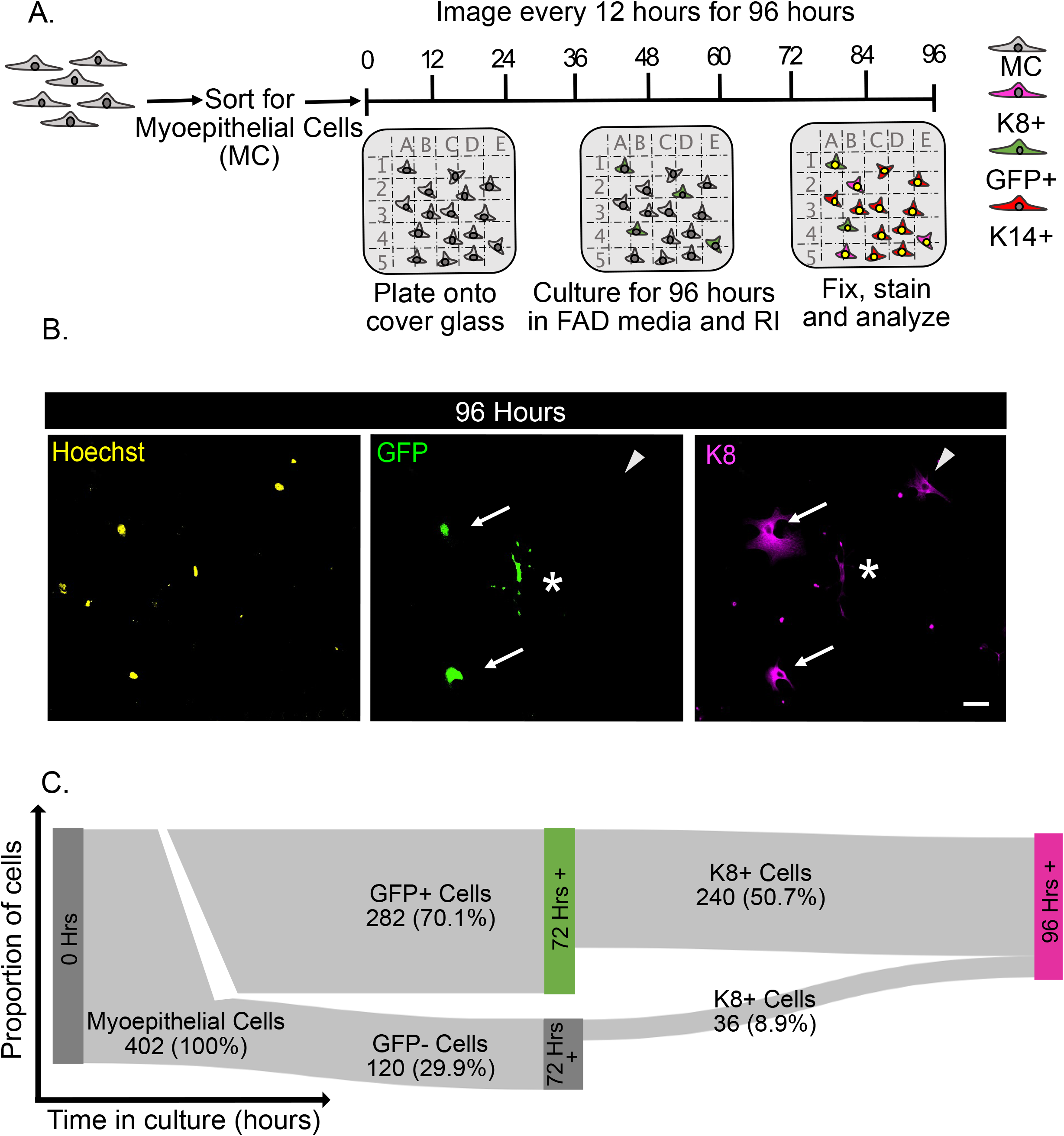
Cell tracking shows that a majority of TG11.5kb myoepithelial cells express GFP prior to expressing luminal cell markers during transdifferentiation. A. Diagram of experimental approach to track transdifferentiation of single myoepithelial cell (MC) conversion in vitro toward the luminal lineage (K8+). B. Representative images of myoepithelial cell transdifferentiation at 96 hrs, and differential conversion paths. Images only showing GFP and K8 expression. Arrows indicate cells that express GFP prior to expressing luminal marker K8. Arrowheads point to cells that express K8 without ever expressing GFP, and the star indicates a cell that becomes GFP+ but did not become K8+. Scale bar = 100 µm. C. Sankey Plot displaying cell gene expression decisions in vitro over time. Majority of cells express the cap cell marker GFP prior to expressing K8 during luminal transdifferentiation.

### RNAseq identifies multiple genes that are differentially expressed between cap cells and mature myoepithelial cells

Myoepithelial cells and GFP+ cap cells were isolated from 5 weeks old female Tg11.5kb-GFP mice and subjected to RNAseq. Principle component analysis confirmed that like-samples had the least variance and clustered together, clearly segregating the two cell types (Supplementary Fig S2 A). We validated the RNAseq data by qRT-PCR of chosen mRNAs that were either less expressed or more highly expressed in cap cells as determined from the RNAseq (Supplementary Fig S2 B). A volcano plot identified multiple genes that were up or down regulated in the GFP+ cap cells versus myoepithelial cells (Fig 3 A). Using a p-value of ≤ 0.05 and a fold change of ≥ 2, 63 genes were significantly upregulated in cap cells and 106 were downregulated, as compared to mature myoepithelial cells. Unsupervised hierarchical clustering also identified a cap cell gene signature corresponding to GO terms associated with organ, tissue and embryonic development, which is consistent with the status of cap cells as progenitors (Fig 3 B, C). TheTg11.5kb-GFP transgenic line drives GFP from an alternate promoter for the SHIP-1 (INPP5D) gene, which expresses a shorter variant of SHIP-1 called s-SHIP (Rohrschneider et al., 2005). Although most of the transcribed s-SHIP sequence is identical to that of full-length SHIP-1, there is a unique 42 nt sequence in the 5’ UTR that differentiates it from the canonical SHIP-1 mRNA sequence. We expected that GFP+ cells would also express this s-SHIP sequence. Indeed, analysis of the RNAseq data identified about 300 (+/- 200 fkpm; n=6) reads of the unique s-SHIP sequence in purified GFP+ cells and zero reads in the mature myoepithelial cells (Fig 3 D).

**Figure 3.**
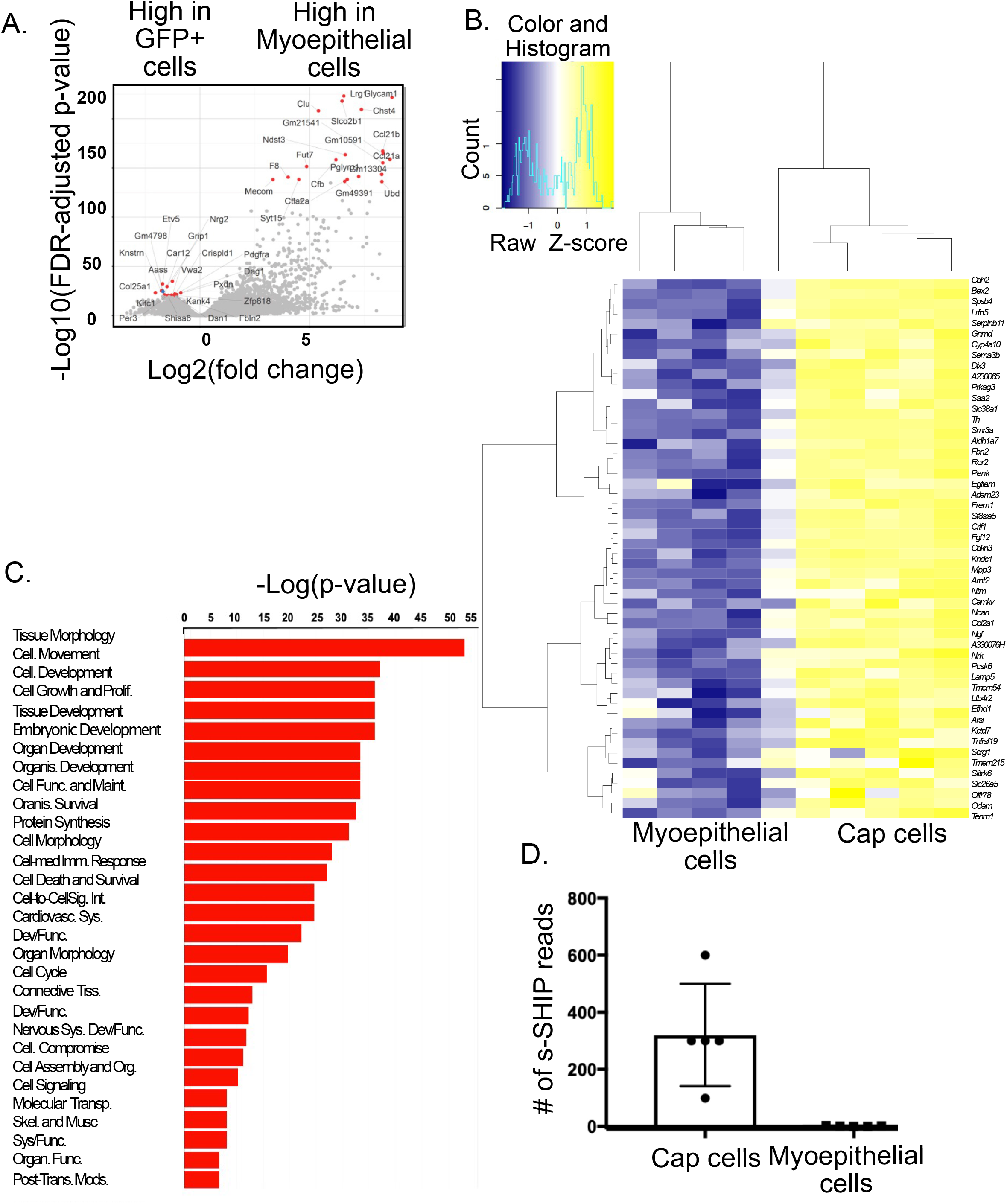
Differential gene expression in myoepithelial cells vs. cap cells. A. Volcano plot displaying differential gene expression with the top 20 most highly expressed genes for each sample in red. Genes filtered with a p-value <0.005. B. Heatmap displaying ∼50 of the top genes differentially expressed between myoepithelial cells and cap cells. Genes filtered with a p-value < .05 and fold change > 2. C. GO ontology graph of biological functions of genes most highly expressed in cap cells. Genes filtered with a p-value of < 0.05 and fold change > 2. D. Dot plot displaying read count values from RNA sequencing data which show that s-SHIP is differentially expressed between GFP+ cap cells and myoepithelial cells. Each dot is a biological replicate. n=5. Error bars = Mean +/- SD.

### Transcription factors predicted to regulate the GFP+ cap cell signature

To determine what transcription factors are regulating the expression of the genes highly expressed in GFP+ cap cells we performed Gene Regulatory Network Analysis using iRegulon, which is a computational method developed to reverse-engineer a transcriptional network underlying a co-expressed gene set using *cis*-regulatory sequence analysis (Janky et al., 2014). This program bases its predictions on databases of ∼10,000 TF motifs and 1000 Chip-Seq data sets. The analysis identified 15 transcription factors predicted to regulate the majority of genes most highly expressed in GFP+ cap cells (Fig 4 A). We determined from the RNAseq data that of these transcription factors RFX3, MTF1, PDX1, and GATA2 are consistently upregulated in cap cells (Fig 4 B).

**Figure 4.**
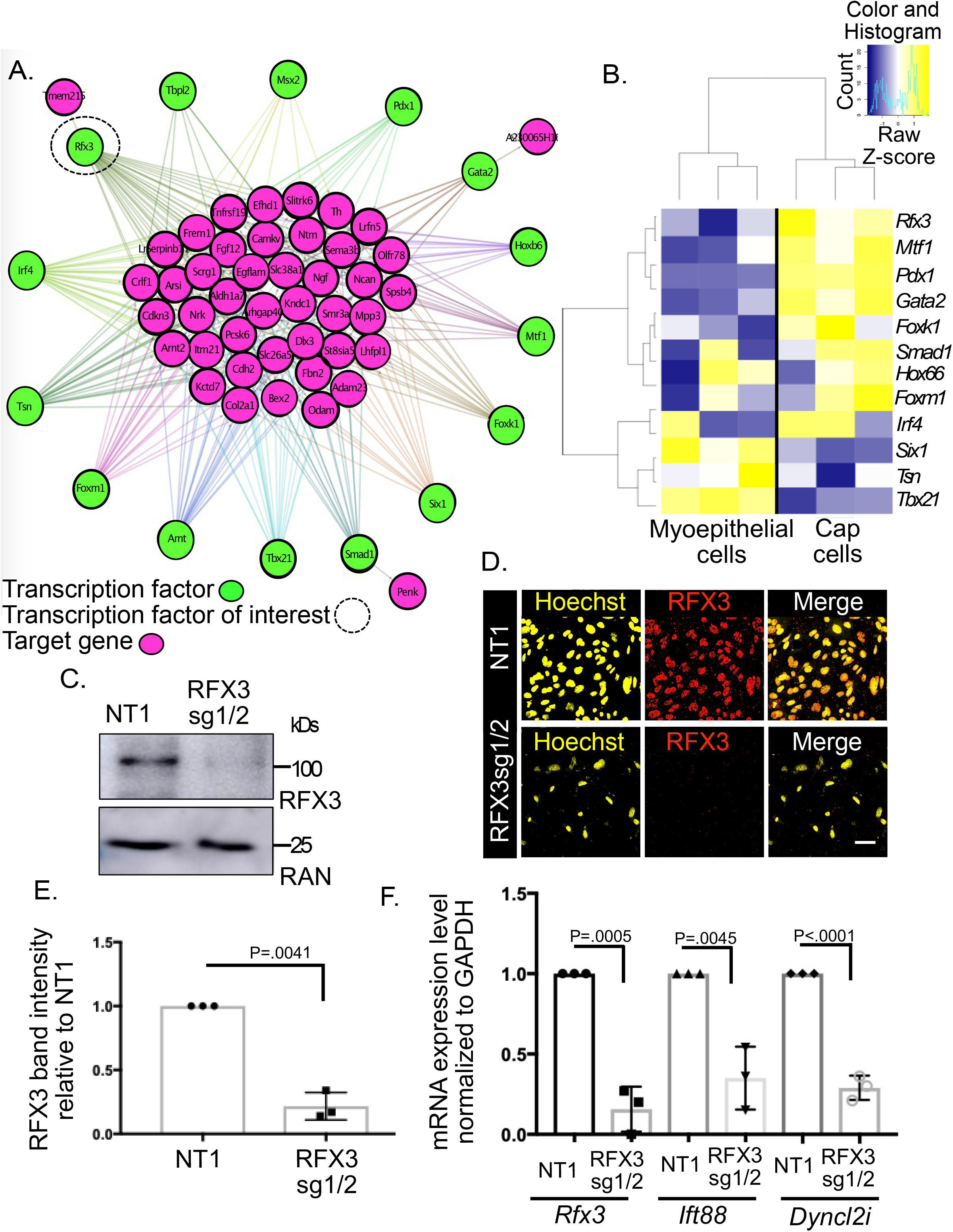
Gene Regulatory Network Analysis identifies Regulatory Factor X3 (RFX3) as a potential upstream regulator of cap cell gene expression. A. iRegulon Web showing the top 15 transcription factors (green) predicted to regulate the differentially expressed genes (DEGs) (pink) most highly expressed cap cells. RFX3 was a transcription factor of interest (black dashed circle). B. Heat map of differentially expressed transcription factors predicted to regulate genes most highly expressed in cap cells. C. Western blot confirming RFX3 KO in commaD Beta cells. anti-RAN is used as the loading control. D. Immunocytochemistry confirming RFX3 knockout (KO) in primary cap cells. NT1 is the non-targeting 1 control lentivector and RFX3sg1/2 are the single guide lentivectors 1 and 2 used to excise the RFX3 DNA binding domain. Scale bar = 100µm F. Quantification of western blot. RFX3 band intensity relative to NT1. P-value = .0041, statistics calculated using unpaired t-test. Error bars = Mean +/- SD. G. qRT-PCR confirms RFX3 loss results in a decrease in target gene (IFT88 and DYNC2LI) expression. P-value = .0005, .0045, <.0001, statistics calculated using unpaired t-test. Error bars = Mean +/- SD.

RFX3 is a key factor in ciliogenesis (Chen et al., 2018). Primary cilia have been reported to regulate branching morphogenesis during mammary gland development (McDermott, Liu, Tlsty, & Pazour, 2010). It was also shown that cilia are enriched in mammospheres and cilia defects decreased the ability to form mammospheres. Loss of IFT88, a target of RFX3, reduces mammosphere growth (Mitchell & Serra, 2014). To determine whether RFX3 is an important regulator of the conversion of myoepithelial cells into GFP+ cap cells, we used Cas9-mediated gene editing with 2 gRNAs to target and splice out the RFX3 DNA binding domain in purified mature myoepithelial cells from the Tg11.5kb-GFP mice. A non-targeting gRNA (NT1) was used as the negative control. A high efficiency of knockout was confirmed on uncloned cultures of the infected cells, as compared to cells transduced with the NT1 gRNA, by immunoblot and immunofluorescence (Fig 4 C,D,E). RT-PCR of RFX3 and of two known targets of RFX3, IFT88 and DYNC2U, were also significantly diminished by the knockout (Fig 4 F). The knockout cells were also grown in culture for 96 hrs then stained for K8. The number of GFP+ cells in the RFX3-culture was significantly lower than in the NT1 culture, while K8+ cell numbers were increased (Fig 5 A,B,C). Moreover, when we performed time-lapse imaging and traced isolated myoepithelial cells back to when they were initially plated, fewer of the RFX3-cells became GFP+ (30% versus 60% for the NT1 control cells), and – surprisingly - more GFP-cells transdifferentiated into the K8+ luminal lineage (Fig 5 D). These data suggest a reduced frequency of de-differentiation towards the cap cell lineage and/or increased transdifferentiation into K8+ luminal cells. We also observed that there was a delay in the onset of GFP expression, and more cells reverted to a non-GFP+ state (Fig 5 E).

**Figure 5.**
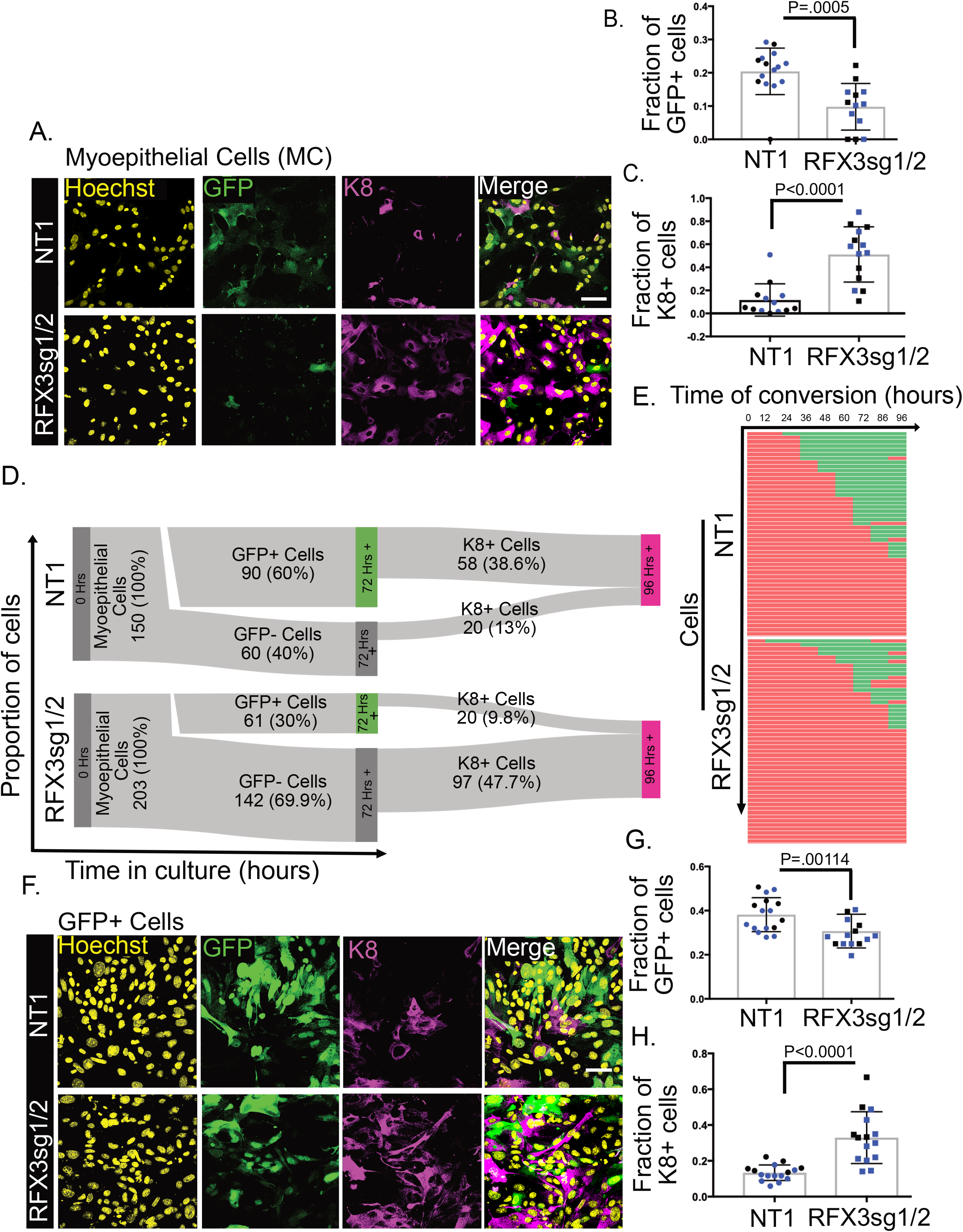
Loss of RFX3 promotes luminal transdifferentiation and reduces GFP+ cap cell abundance. A. Representative images of RFX3 knockout (RFX3sg1/2) versus a negative control (non-targeting gRNA) in myoepithelial cells cultured in vitro for 96 hrs. Scale bar = 100µm B. Quantification showing a decrease in the number of myoepithelial cells that become GFP+ (NT1: n= 14, 2 experiments, RFX3sg1/2: n=13, 2 experiments) when RFX3 is lost. P-value = .0005, statistics calculated using mixed model ANOVA. Error bars = Mean +/- SD. C. Quantification graph showing an increase in the fraction of luminal cells (K8+) (NT1:n= 13, 2 experiments, RFX3sg1/2:n=14, 2 experiments) that arise. Individual data points show values for technical replicates; different colors identify biological replicates. Statistics calculated using mixed model ANOVA. Error bars = Mean +/- SD. D. Sankey Plots of NT1 and RFX3 knockout (RFX3sg1/2) transdifferentiation behavior in vitro after 96 hrs. Loss of RFX3 results in a majority of myoepithelial cells converting towards a luminal cell fate, bypassing a GFP+ phase. E. Kymograph showing times of conversion to GFP expression of myoepithelial cells. n= 50 cells F. Representative images of RFX3 knockout (RFX3sg1/2) or the non-targeting control in FACS-purified GFP+ cap cells cultured in vitro for 96 hrs. Scale bar = 100µm G. Quantification showing a decrease in the number of myoepithelial cells that become GFP+ (NT1: n= 16, 2 experiments, RFX3sg1/2: n=14, 2 experiments) when RFX3 is lost. Individual data points show values for technical replicates; different colors identify biological replicates. Statistics calculated using mixed model ANOVA. Error bars = Mean +/- SD. H. Quantification graph showing an increase in the number of luminal cells (NT1: n= 16, 2 experiments, RFX3sg1/2: n=15, 2 experiments) that arise. P-value < 0.0001, statistics calculated using mixed model ANOVA. Error bars = Mean +/- SD.

To further investigate mechanism, we repeated the experiment using RFX3 knockout in purified GFP+ cap cells. As shown in Fig 5 F,G,H the loss of RFX3 reduced the GFP+ population and increased the fraction of K8+ cells. This result suggests that RFX3 normally functions to stabilize cap cell identity and prevent transdifferentiation into the luminal lineage. Because luminal cells are not competent to regenerate mammary glands, a prediction of this model is that deletion of RFX3 would reduce the capacity of basal cells (cap or myoepithelial cells) to form mammospheres, or to regenerate mammary glands in vivo after transplantation into the cleared fat pads of recipient mice. Indeed, we observed a significant reduction in mammosphere growth of myoepithelial cells grown in 3D culture (Fig 6 C,D). Importantly, this effect was not the result of a decreased ability of RFX3 knockout cells to proliferate, as we could detect no significant difference in BrdU incorporation into these cells versus an NT1 control that expresses a non-targeting gRNA (Fig 6 A,B).

**Figure 6.**
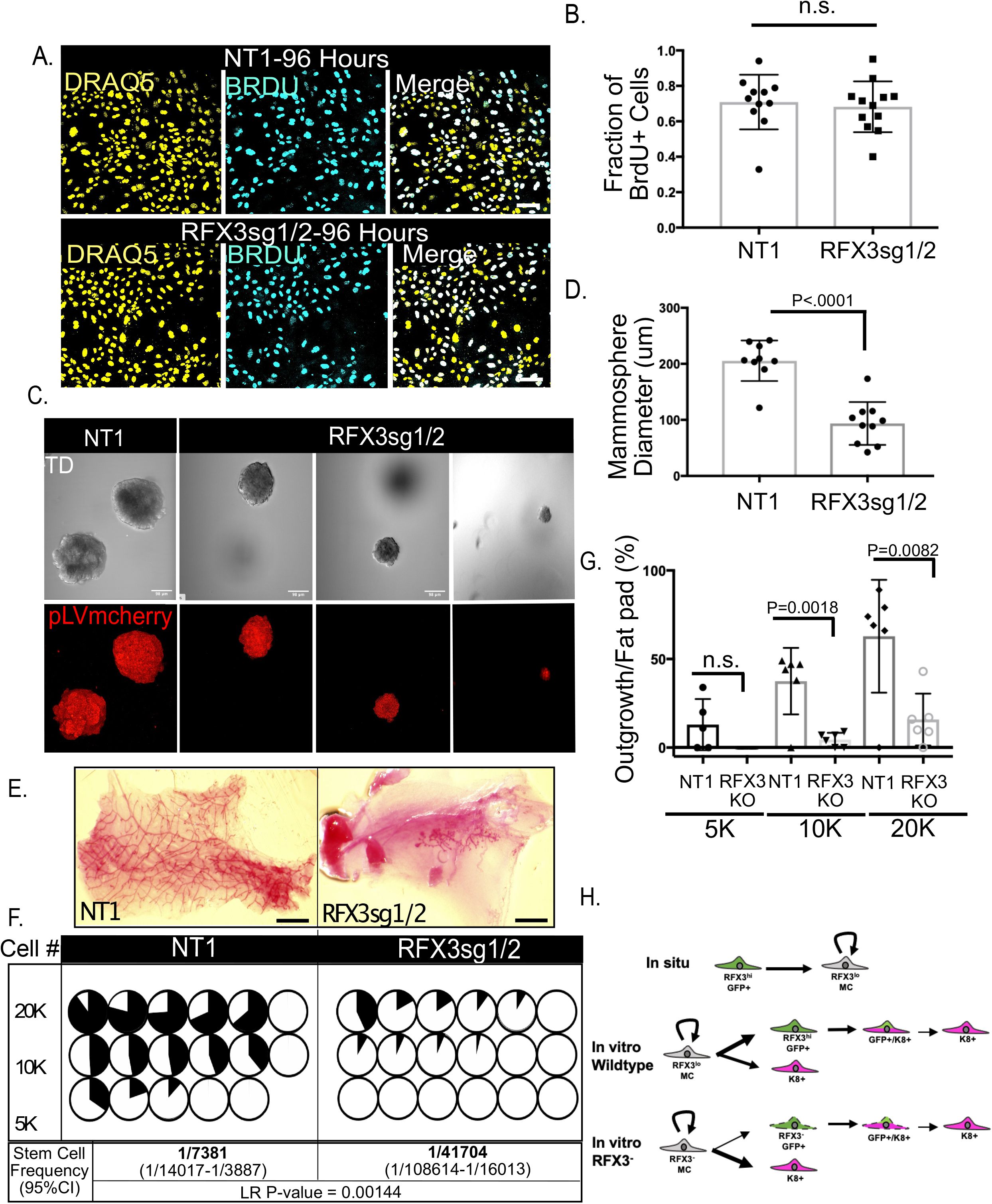
RFX3 stabilizes basal cell identities. A. Representative images of myoepithelial cells transduced with either NT1 or RFX3 knockout lentivirus (RFX3sg1/2), pulse-labeled for 2 hrs with BrdU and stained for DNA (DRAQ5) and BrdU incorporation. Scale bar = 100µm. B. Quantification of fraction of BrdU+ myoepithelial cells. P-value= n.s., calculated using unpaired t-test. Error bars = Mean +/- SD. C. Representative images of myoepithelial cells transduced with either NT1 or RFX3 knockout lentivirus (RFX3sg1/2) and grown as mammospheres. Cherry+ mammospheres confirm transduction. Scale bar = 98µm D. Quantification of mammosphere size. P-value = < 0.0001, calculated using unpaired t-test. Error bars = Mean +/- SD. E. Representative images of outgrowths regenerated from transplantation of 20K NT1- or 20K RFX3sg1/2–transduced myoepithelial cells. Scale bar = 3 mm. F. Circle graphs displaying the percentage of outgrowth that resulted (outgrowth/fat pad area) from the number of cells transplanted. Stem cell frequency was calculated and generated a likelihood ratio (LR) of P-value = 0.00144. G. Quantification of transplantation assay outgrowth. P-values were calculated using unpaired t-test. P-value = n.s. (5K cells) 0.0018 (10K cells), 0.0082 (20K cells). Error bars = Mean +/- SD. H. Model for the role of RFX3 in stabilizing mammary basal cell identities. In situ, cap cells, the myoepithelial progenitors (GFP+), give rise only to mature myoepithelial cells (MC) which maintain their lineage through self-renewal. In vitro, wildtype myoepithelial cells de-differentiate predominantly towards a progenitor state (GFP+) before changing lineage into K8+ luminal cells (K8). A subset of myoepithelial cells can apparently bypass the GFP+ state and transdifferentiate directly into K8+ cells. RFX3 KO myoepithelial cells have a reduced frequency of conversion into a GFP+ state (thin arrow) and increased transdifferentiation directly into a luminal cell lineage (thick arrow).

We next tested the efficiency of mammary gland regeneration in a transplantation assay. Five, ten or twenty thousand unsorted mammary epithelial cells were transplanted into the cleared fat pads of isogenic recipient mice. After 8 weeks the mice were euthanized and the glands were removed, fixed, and stained with Carmine Alum. Analysis of these whole mounts showed that mammary gland outgrowth was substantially reduced for transplanted RFX3-negative cells compared to the NT1 control (Fig 6 E,F,G). We infer that the RFX3 transcription factor functions to stabilize the identities of basal cell populations, which is necessary for efficient ductal regeneration.

## DISCUSSION

It has become clear in recent years that the concept of distinct, stable cell states that change in a uni-directional manner during development is incorrect, and that most cells in most animal tissues are in metastable states that – often in response to stress – can revert to earlier states or change lineages. For example, following injury, differentiated airway epithelial cells and intestinal epithelial Dll1+ secretory progenitors can become functional stem cells (Tata et al., 2013; van Es et al., 2012). Injury to the heart in zebrafish results in dedifferentiation and proliferation of cardiomyocytes (Jopling et al., 2010). Differentiated cells can also be forced to revert to a stem cell state by the over-expression of various transcription factors, most famously for mesenchymal fibroblasts, which in the context of the Yamanaka factors are reprogrammed into pluripotent stem cells. Mature pancreatic acinar cells can be converted directly into beta cells by expression of Ngn3, Pdx1 and Mafa transcription factors (Zhou, Brown, Kanarek, Rajagopal, & Melton, 2008); and mammary luminal cells can be driven to a multipotent stem-like cells by expression of Sox9 and Slug (Guo et al., 2012). On the other hand, mature mammary myoepithelial cells, which in vivo normally form a discrete, self-renewing population with no evidence of multipotency, can spontaneously convert to a stem-like state after isolation from the mammary gland and grown in culture, or transplanted into a recipient fat pad (Prater et al., 2014; Van Keymeulen et al., 2011). This conversion suggests that the micro-environment plays an important role in stabilizing the differentiated state of mature myoepithelial cells, a hypothesis consistent with recent demonstrations that DNA damage to the mammary gland or the ablation of luminal cells by diphtheria toxin in vivo results in transdifferentiation of myoepithelial cells to the luminal lineage. However, the mechanism for this conversion remains unclear. Do myoepithelial cells directly switch lineage, or do they pass through an intermediate, stem-like state? And what gene regulatory network controls lineage stability versus instability?

To address these questions, we examined the transdifferentiation of mature mammary myoepithelial cells towards the luminal lineage when they are isolated from the mammary gland and grown in culture. Single cell RNAseq showed that over a period of 96 hrs, a fraction of the myoepithelial cells take on a luminal identity in vitro, and cell tracking showed that the majority of such cells pass through an intermediate state in which they express a marker of cap cells, which are myoepithelial cell progenitors. Differential gene expression analysis and regulatory network analysis using iRegulon identified several transcription factors that potentially control expression of multiple genes that are upregulated in cap cells. One of these, RFX3, proved to be necessary for the stability of basal (cap and myoepithelial) cell identities. Knockout of RFX3 strongly reduced the number of cells that expressed the cap cell GFP marker. Moreover, while loss of RFX3 had no impact on proliferation of myoepithelial cells in vitro, it significantly reduced mammosphere growth and reduced mammary ductal outgrowth in a transplantation assay. These data suggest that RFX3 is not required for cap cell identity, or for conversion of myoepithelial cells to the luminal lineage, but instead plays a role in stabilizing cell identity. We speculate that state stabilization is an important process in development, and that multiple transcription factors, regulated by signaling from the micro-environment, might operate in different contexts and tissues to ensure phenotypic stability.

## ACKNOWLEDGEMENTS

This work was supported by grant R35 CA197571 to IGM, and T32 HD07502 to EMT. We thank members of the Macara laboratory for advice and support.

## RESOURCE AVAILABILITY

### Lead Contact

Further information and requests for resources and reagents should be directed to and will be fulfilled by the lead contact, Ian Macara, (615) 875-5565.

### Materials Availability

This study generated Rfx3 sgRNAs (KO 1: ATCATGCAGACTTCAGAGA, KO 2: GCAAGTGCCAGTGCAGCAGC) within the pLentiCRISPR-mCherry (Addgene plasmid #75161;http://n2t.net/addgene:75161;RRID:Addgene_75161) backbone that target the Mus musculus Rfx3 (Gene ID: 19726) DNA-binding domain.

### Data and Code Availability

The RNA sequencing datasets generated in this study will be deposited to the GEO repository on the NCBI website.

## KEY RESOURCES TABLE

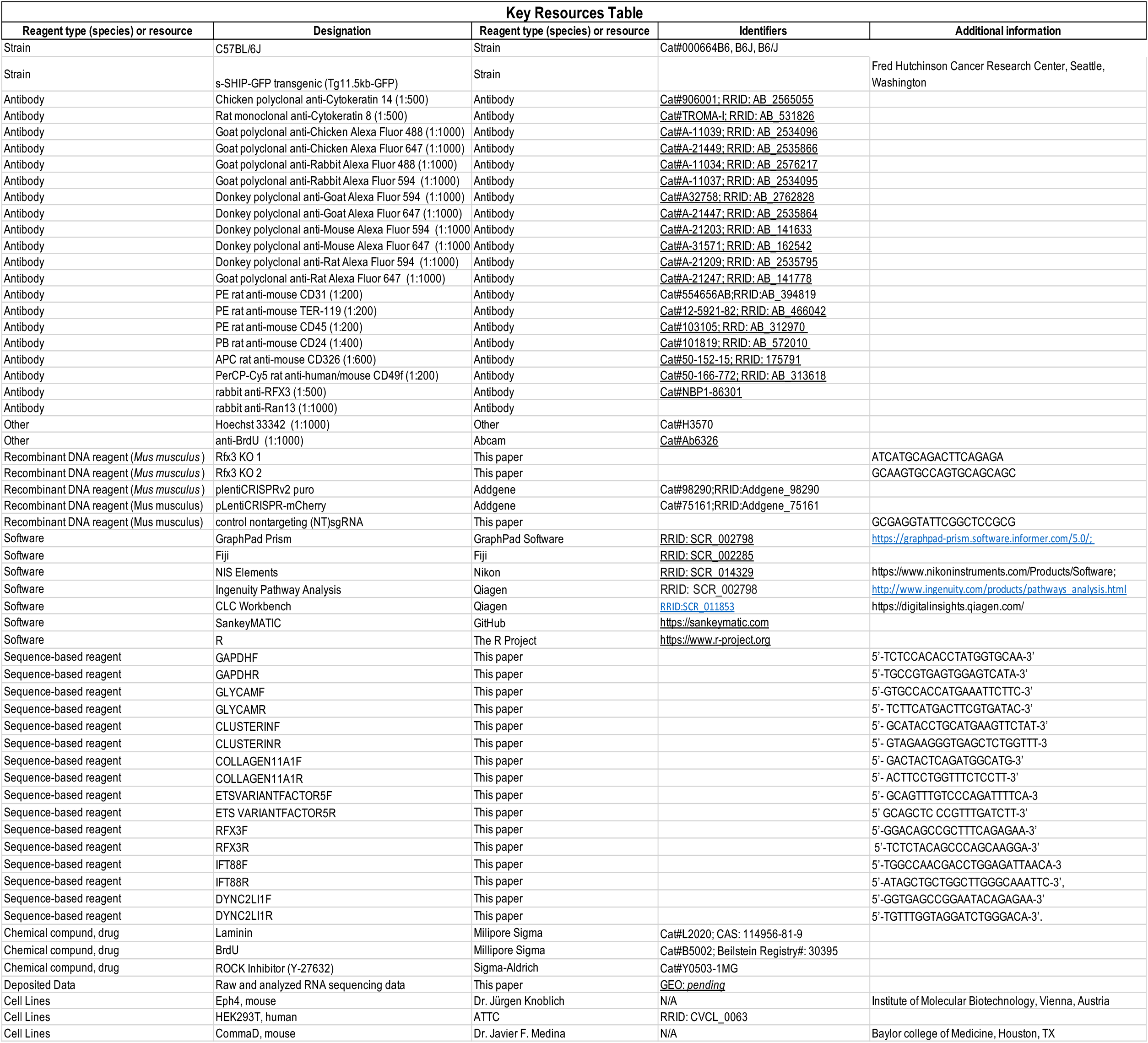

## MATERIALS AND METHODS

### Mice

C57BL/6J (JAX stock # 000664) female mice were purchased from Jackson Laboratory (Bar Harbor, ME). The s-SHIP–GFP transgenic (Tg11.5kb–GFP) mice were provided by Dr. Larry Rohrschneider from Fred Hutchinson Cancer Research Center, Seattle, Washington. Mice were housed in the Vanderbilt mouse facility with a standard 12hrs light/12hrs dark cycle and provided with normal laboratory chow and water. The mice were monitored daily by the Vanderbilt Division of Animal Care (DAC). s-SHIP-GFP primer sequences used for genotyping were as described previously (Bai and Rohrschneider, 2010). All mouse experiments were performed with approval from the Vanderbilt Institutional Animal Care and Use Committee (IACUC).

### Cell Lines

Mouse mammary EpH4 cells were provided by Dr. Juergen Knoblich (Institute of Molecular Biotechnology, Vienna, Austria). HEK293T (ATCC CRL-3216) cells were obtained from ATCC. CommaDβ cells were provided by Dr. Medina (Baylor college of Medicine, Houston, TX). Cell lines were cultured in Dulbecco’s modified Eagle medium (Life Technologies), supplemented with 10% Fetal Bovine Serum (R&D Systems, Minneapolis, MN) and 1X penicillin/streptomycin (Life Technologies) and maintained in culture at 37°C with 5% CO_2_. WPI, psPAX2 and pMD2.G were provided by Didier Tron. LentiCRISPRv2 puro was a gift from Brett Stringer (Addgene plasmid # 98290; http://n2t.net/addgene:98290;RRID:Addgene_98290). pLentiCRISPR-mCherry was a gift from Beat Bornhauser. (Addgene plasmid #75161;http://n2t.net/addgene:75161;RRID:Addgene_75161). Guide RNAs for RFX3 DNA binding-domain knock out (Key Resources Table) were cloned into plentiCRISPRv2 puro or pLentiCRISPR-mCherry. sgRNAs were ligated into lentivectors at the *BsmBI* restriction site as described in the Zhang lab protocol (Sanjana, Shalem, & Zhang, 2014; Shalem et al., 2014).

### Lentiviral Production

Lentivirus was produced by transfecting 80% confluent 15 cm dishes of HEK293T cells. Lentiviral plasmid, protein plasmid (pMD2.G) and packaging plasmid (psPAX2) were transfected into HEK293T cells at a 1.5:1:1 ratio using calcium phosphate precipitation. Virus-conditioned medium was collected from cells 48 hrs post transfection, stored at −80 °C, and titered using HEK293Tcells.

### Primary Cell Isolation

To isolate mammary epithelial cells the third and fourth pairs of mammary glands were removed from 4 to 6-week-old s-SHIP–GFP transgenic (Tg11.5kb–GFP) mice, minced with scissors and digested in digestion medium (DMEM/F12, 2 mg ml^−1^collagenase I (Roche), 5 mg ml^−1^ insulin (Sigma), 100 U ml^−1^ penicillin/streptomycin) for 1 hr at 37 °C. Epithelial organoids were collected by centrifugation at 1500 rpm for 5 min. The cell pellet was washed for 1 min in 5 ml of DMEM/F12 containing DNase I and centrifuged at 1500 rpm for 5 min. The pellet was resuspended in 5 ml DMEM/F12 10% fetal bovine serum followed by 15 sec of centrifugation at 1500 rpm five times. Cells were resuspended in 1 ml of fresh trypsin/EDTA (Invitrogen) and incubated at 37 °C for 15 min, dissociated into a single-cell suspension and passed through a cell strainer (BD) to obtain a single-cell suspension of mammary gland cells. Primary cells were transduced with lentivector (RFX3 KO lentivectors or NT1) by incubating for 1-4 hours at 37 °C.

### BrdU Incorporation Assay

72 hours post-transduction with lentivector (RFX3 KO or NT1), myoepithelial cells were pulse-labeled for 2 hrs with BrdU (Millipore Sigma, Cat # L2020). Cells were then stained with anti-BrdU (Abcam, Cat# Ab6326) according to the antibody protocol.

### Flow cytometry, antibodies, and cell sorting

Single mammary epithelial cells were blocked in 5% goat serum for 5 min on ice, stained with antibodies for 30 min on ice, washed, and resuspended in 1X PBS, 2mM EDTA, 2%FBS. Antibodies used were PE rat anti-mouse CD31 (1:200; BD Biosciences), PE rat anti-mouse TER-119 (1:200; Invitrogen), PE rat anti-mouse CD45 (1:200; BioLegend), PB rat anti-mouse CD24 (1:400; BioLegend), APC rat anti-mouse CD326 (Epcam) (1:600; Ebioscience), and PerCP-Cy5 rat anti-human/mouse CD49f (1:200; BioLegend). The single live cells were gated and sorted on 5-laser FACS Arialll flow cytometers.

### Myoepithelial Cell Conversion Assays

Freshly sorted mature myoepithelial cells were plated on 16-well chambered coverglasses (Grace Biol-Labs Cat. # 112358) in FAD media supplemented with 10 μm ROCK Inhibitor (Y-27632) (Sigma-Aldrich Cat.# Y0503-1MG) and cultivated for 96 hrs. Media was changed every day and cells were fixed, stained and analyzed at 96 hrs.

### Cell Fate Mapping

Myoepithelial cells were sparsely plated onto gridded coverslips (ibidi catalogue #80826-G500) in FAD media and live imaged on a Nikon A1R inverted confocal microscope (Nikon Instruments Inc.) every 12 hrs for 96 hrs using 20X/0.75 NA. At 96 hrs cells were fixed and stained for K8 and nuclei. Cells were annotated at 24 hrs and tracked up to hr 96. Cells that were tracked over the 96 hrs were scored for becoming GFP+ and whether the cells were expressing luminal marker (K8) at hr 96. Sankey plots were generated using SankeyMATIC by Steve Bogart (https://github.com/nowthis/sankeymatic/blob/main/README.md). Kymographs were generated by analyzing images from 12 hrs to 96 hrs and marking the timepoint at which GFP expression is visible and the timepoint when GFP expression turns off, if within the 96 hrs.

### Mammosphere Assays

Myoepithelial cells were embedded in growth-factor-reduced Matrigel (BD) on16-well chambered coverglasses (Grace Biol-Labs Cat. # 112358) and cultured in mammosphere medium supplemented with 10 μm Y-27632 and cultivated for 10 days. Medium was changed every day and cells were fixed, stained and analyzed on day 10.

### Tissue Processing, Staining and Analysis

For immunocytochemistry, cells were fixed in 4% paraformaldehyde at room temperature for 15 min. Cells were permeabilized with 0.2% Triton X-100. For immunohistochemical analyses mammary tissue samples were cryo-embedded in O.C.T. (Fisher Scientific, Hampton, NH) and cryo-sectioned on a Leica C1950 cryostat generating 50 μm sections of mammary tissue. Tissue sections were fixed for 15 min in 4% paraformaldehyde or for 5 min in -20°C Acetone. Both cells and tissues were blocked in 1x Western Blocking Reagent (Roche). The primary antibodies used in this study included: chicken α-cytokeratin 14, rabbit α-S100A4 (BioLegend, San Diego, CA), rat α-cytokeratin 8 (Developmental Studies Hybridoma Bank, Iowa City, Iowa), rabbit α-rfx3 (Novus Biologicals, Centennial, CO). Secondary antibodies used in this study included: Alexa Fluor 405, 488, 594 and 647 conjugates of anti-chicken, -mouse, -rat, -rabbit and -goat (ThermoFisher Scientific, Waltham, MA). Hoechst 33342 (ThermoFisher Scientific, Waltham, MA) was used to stain DNA. Slides were mounted in Fluoromount G and sealed with nail varnish. Laser scanning confocal Images were acquired on a Nikon A1R inverted confocal microscope (Nikon Instruments Inc.). 20X/0.75 numerical aperture (NA) and 40X/1.30 NA Plan Apochromat objectives were use. Type B Immersion oil (Cargille Laboratories, Cedar Grove, NJ) was used. Image post-acquisition processing was done using Nikon NIS-Elements imaging software and Fiji (ImageJ) software. The images in the manuscript are maximum intensity projections of z-stacks and merged images are composites of individual color channels.

### RNA Sequencing

Total RNA (50-200ng) isolation was performed using either TRIzol (Life Technologies) or the RNeasy Mini Kit (Qiagen, Hilden, Germany). 5 samples of myoepithelial cells and 5 samples of s-SHIP GFP+ cap cells were submitted and subjected to quality analysis using a 2100 Bioanalyzer (Agilent Technologies, Santa Clara, CA). All Samples had an average RNA integrity number (RIN) value ∼8. Libraries for whole transcriptome analysis were generated following Illumina’s TruSeq RNA v2 sample preparation protocol. Libraries were sequenced on an Illumina HiSeq 3000 at the Vanderbilt Technologies for Advanced Genomics (VANTAGE). 150 million reads at 75 basepairs were obtained for each of the samples. Data processing, analysis and plotting were performed using R software, CLC Genomics Workbench and Ingenuity Pathway Analysis (Qiagen, Hilden, Germany). Heatmaps were generated using R graphics on RNAseq gene lists filtered with a p-value < .05 and fold change >2. Volcano plots were generated using CLC workbench with genes filtered with a p-value of <0.005. The GO ontology graph was generated using Ingenuity Pathway Analysis and genes were filtered with a p-value of <.05 and fold change >2. Reads for s-Ship were quantified manually by analyzing the read alignment data to the 42 nucleotide unique sequence present in this variant.

### Single Cell RNA Sequencing

After 96 hrs cultivation, converted myoepitheial cells were dissociated using TrypLE and cell viability was determined using Trypan Blue. The control sample (fresh s-SHIP GFP+ cap cells, myoepithelial cells and luminal cells) and cultivated cells were encapsulated and barcoded using the inDrop platform (1CellBio) with an in vitro transcription library preparation protocol (Klein et al., 2015). As per Klein et al., the CEL-Seq workflow is summarized: 1) RT, 2) ExoI, 3) SPRI purification (SPRIP), 4) SSS, 5) SPRIP, 6) T7in vitro transcription linear Amplification, 7) SPRIP, 8) RNA Fragmentation, 9) SPRIP, 10) primer ligation, 11) RT, 12) library enrichment PCR. The number of cells encapsulated was calculated by the density of cells arriving at the device multiplied by the duration of encapsulation. After library preparation, the samples were sequenced using Nextseq 500 (Illumina) using a 150bp paired-end sequencing kit. After sequencing, reads were filtered, sorted by their barcode of origin and aligned to the reference transcriptome using the inDrops pipeline (https://github.com/indrops/indrops). Mapped reads were quantified into UMI-filtered counts per gene, and barcodes that correspond to cells were retrieved based on previously established methods (Klein et al., 2015).

### Real-time qPCR

cDNA was reverse transcribed using the SuperScript III First-Strand Synthesis System (Invitrogen). qPCR was performed with triplicate replicates on a BioRad CFX96 Thermocycler and analyzed using the ΔΔCt method. Expression levels were calculated relative to GAPDH. Primer sequences used were: GAPDHF 5’-TCTCCACACCTATGGTGCAA-3’, GAPDHR 5’-TGCCGTGAGTGGAGTCATA-3’, GLYCAMF 5’-GTGCCACCATGAAATTCTTC-3’, GLYCAMR 5’-TCTTCATGACTTCGTGATAC-3’, CLUSTERINF 5’-GCATACCTGCATGAAGTTCTAT-3’, CLUSTERINR 5’-GTAGAAGGGTGAGCTCTGGTTT-3’, COLLAGEN11A1F 5’-GACTACTCAGATGGCATG-3’, COLLAGEN11A1R 5’-ACTTCCTGGTTTCTCCTT-3’, ETSVARIANTFACTOR5F 5’-GCAGTTTGTCCCAGATTTTCA-3’, ETS VARIANTFACTOR5R 5’ GCAGCTC CCGTTTGATCTT-3’,RFX3F 5’-GGACAGCCGCTTTCAGAGAA-3’, RFX3R 5’-TCTCTACAGCCCAGCAAGGA-3’,IFT88F 5’-TGGCCAACGACCTGGAGATTAACA-3’,IFT88R 5’-ATAGCTGCTGGCTTGGGCAAATTC-3’, DYNC2LI1F 5’-GGTGAGCCGGAATACAGAGAA-3’, DYNC2LI1R 5’-TGTTTGGTAGGATCTGGGACA-3’.

### Gene Network Regulatory Analysis

iRegulon (Janky et.al., 2014) was used to predict transcription factor regulation. RNA seq data was filtered for genes highly expressed in GFP+ cap cells with a p-value < .05. Prediction data was generated by surveying transcription factor binding motifs present 20 kb around transcription start site (TSS) [TSS−10 kb, TSS+10 kb]. Gene regulatory networks were generated within Cytoscape.

### Transplantation/ Limited Dilution Assays

Myoepithelial cells were FACS sorted and transduced with either non-targeting or RFX3-targeting lentivirus. 24 hrs later cells were dissociated in trypLE Select (Gibco) for 8 minutes. Dilutions of 5, 000, 10,000, or 20,000 cells were resuspended in 10 ul of injection medium (10 μm Y-27632 (Sigma-Aldrich Cat.# Y0503-1MG), 20% Matrigel (Corning), DMEM/F12, 40 ng/mL EGF, 20 ng/mL (R&D Systems), FGF2 (R&D Systems) and injected into the cleared fat pads of 3-week-old female C57Bl6 mice (Jackson Laboratories) using a 26 gauge needle and Hamilton syringe. Mice were sacrificed 8 weeks post transplantation and whole mounts analyzed. Outgrowths were detected by carmine alum whole-mount. Pictures were acquired with an Olympus SZX16. A positive outgrowth was described as 10% or greater percentage of outgrowth/fatpad.

### Western Blots

Cells were washed with 1x PBS and lysed in 1 ml of lysis buffer containing 20 mM HEPES; pH 7.4, 50 mM NaCl, 2 mM EDTA, and 0.1% Triton X-100, supplemented with cOmplete™mini EDTA-free protease inhibitor cocktails (Roche) and PhosStop (Roche). Cell lysates were briefly centrifuged at 16,000 x g and the soluble fraction was boiled with SDS sample buffer for 5 min. Antibodies used for Western blotting include: rabbit anti-RFX3 (Novus Biologicals, Centennial, CO) and rabbit anti-Ran13 (Macara Lab; (Richards, Lounsbury, & Macara, 1995).

### Statistics

All cell counts were performed using Hoechst staining and a fluorescent antigen marker in NIS-Elements imaging software or Fiji (ImageJ) software. All measurements for mammosphere diameter, transplant outgrowth area and western blot analysis were done using ImageJ. All statistical analyses were performed using unpaired Student’s t-test or two-way ANOVA test. Stem cell frequencies were calculated using Extreme Limiting Dilution Analysis (ELDA) (Hu & Smyth, 2009).

