## Supplementary Figures for "Transcription Factor RFX3 Stabilizes Mammary Basal Cell Identity"

### SUPPLEMENTARY FIGURES AND FIGURE LEGENDS

Supplementary Figure 1.

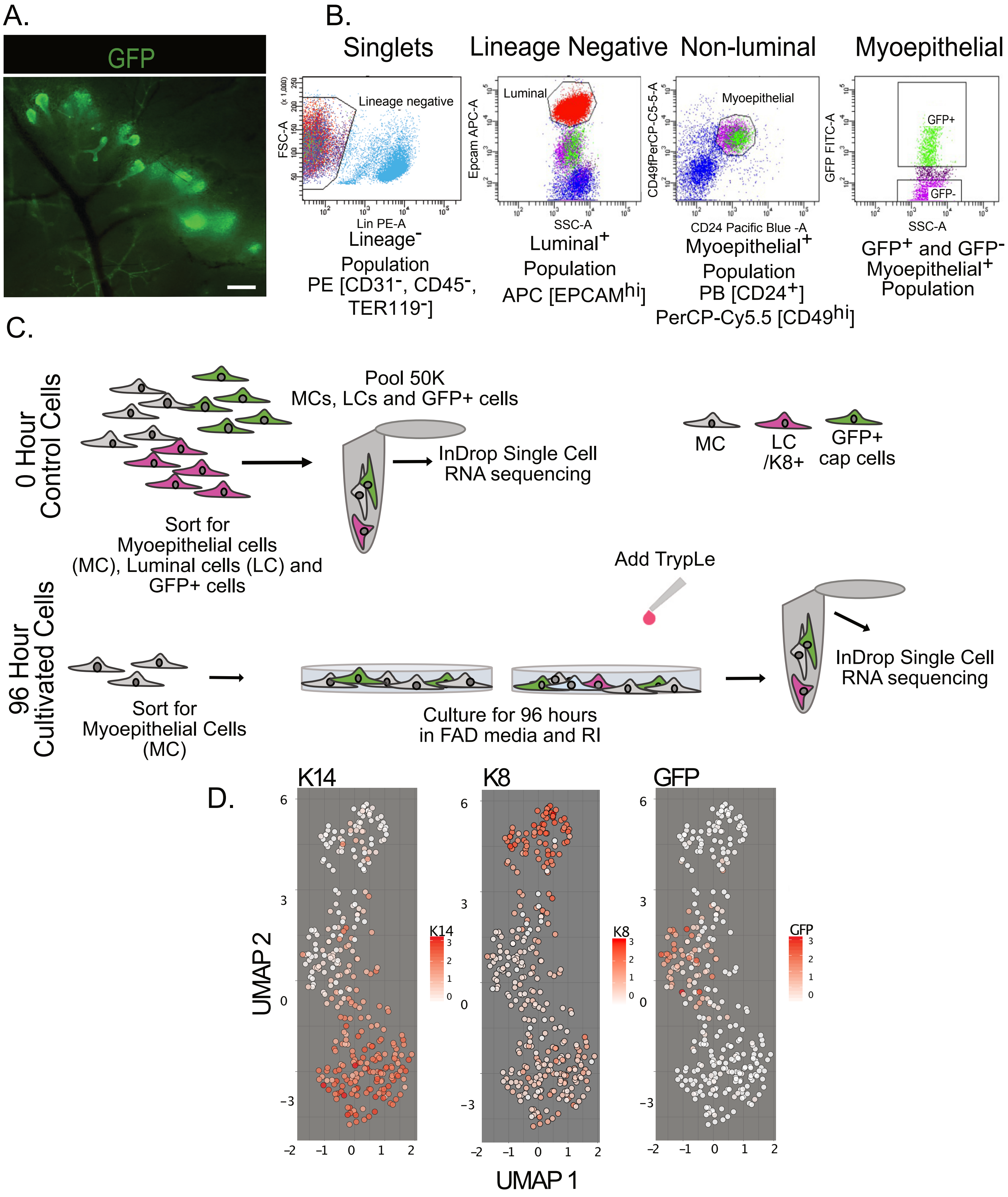

**Supplemental Figure 1. Identification, isolation and single-cell RNA sequencing of GFP+ cap cells and myoepithelial cells**

A. Whole mount of mammary glands from transgenic mice (TG11.5kb-GFP) showing GFP+ terminal end buds. Scale bar = 1 mm
B. Gating strategy for isolation of myoepithelial cells (GFP-) from TG11.5kb-GFP transgenic mice.
C. Diagram of experimental approach to prepare cells for single-cell RNA sequencing.
D. UMAP projections of control sample displaying the 3 sorted and distinctly clustered cell populations with K14 (myoepithelial cells), K8 (luminal cells) and GFP (cap cell) gene expression levels.

Supplementary Figure 2.

A.

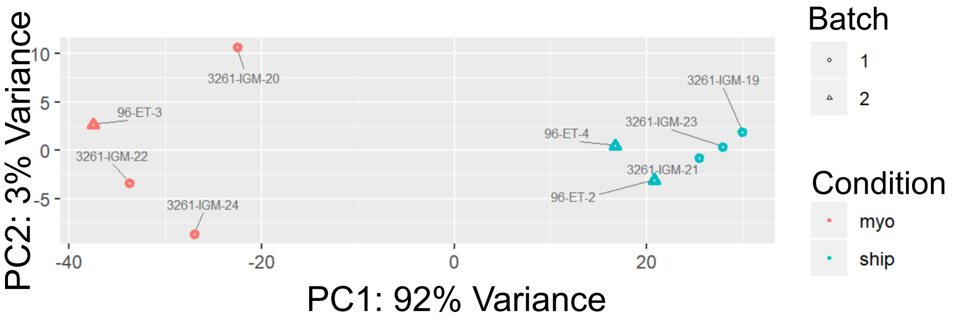


B.

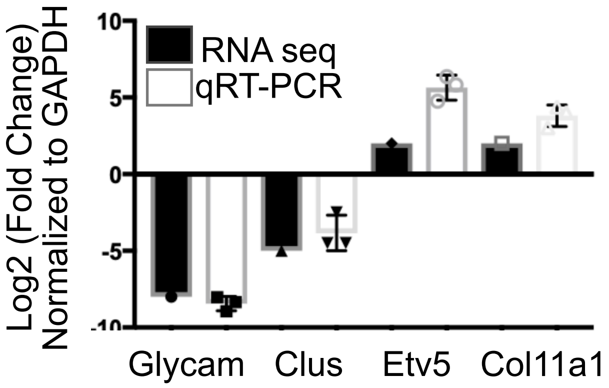

**Supplemental Figure 2. Bulk RNA sequencing validation**

A. Principle Component Analysis of GFP+ cap (ship) and myoepithelial (myo) cells show distinct clustering (data has been batch corrected). Shapes indicate different experiments. ship, n=5, myo, n=4.

B. Validation of RNA seq differential gene expression by qRTPCR. Comparison of the Log2 (Fold Change) (mean) of RNA Seq and qRT-PCR data expression of myoepithelial cells vs. cap cells, which are consistent with one another, n=3. Error bars = mean +/- SD
